# Loter: A software package to infer local ancestry for a wide range of species

**DOI:** 10.1101/213728

**Authors:** Thomas Dias-Alves, Julien Mairal, Michael G.B. Blum

## Abstract

Admixture between populations provides opportunity to study biological adaptation and phenotypic variation. Admixture studies rely on local ancestry inference for admixed individuals, which consists of computing at each locus the number of copies that originate from ancestral source populations. Existing software packages for local ancestry inference are tuned to provide accurate results on human data and recent admixture events. Here, we introduce Loter, an open-source software package that does not require any biological parameter besides haplotype data in order to make local ancestry inference available for a wide range of species. Using simulations, we compare the performance of Loter to HAPMIX, LAMP-LD, and RFMix. HAPMIX is the only software severely impacted by imperfect haplotype reconstruction. Loter is the less impacted software by increasing admixture time when considering simulated and admixed human genotypes. For simulations of admixed Populus genotypes, Loter and LAMP-LD are robust to increasing admixture times by contrast to RFMix. When comparing length of reconstructed and true ancestry tracts, Loter and LAMP-LD provide results whose accuracy is again more robust than RFMix to increasing admixture times. We apply Loter to individuals resulting from admixture between *Populus trichocarpa* and *Populus balsamifera* and lengths of ancestry tracts indicate that admixture took place around 100 generations ago. We expect that providing a rapid and parameter-free software for local ancestry inference will make more accessible genomic studies about admixture processes.

## Introduction

Admixture or hybridization between populations is a natural phenomenon that provides opportunity to map genomic regions involved in phenotypic variation and biological adaptation (Buerkle and Lexer 2008; Payseur and Rieseberg 2016). Mapping can rely on Local Ancestry Inference (LAI) of admixed individuals, which consists of computing at a given locus the number of copies that originates from the ancestral source populations. LAI uses haplotypes from putative source populations and processes haplotypes or genotypes from admixed population to infer local ancestry of admixed individuals. Figure 1 shows local ancestry of four simulated Populus individuals resulting from admixture between two Populus species (Suarez-Gonzalez et al. 2016). Sequence and dense genotype data are now generated for a wide range of species besides humans for which LAI is relevant. LAI can be used to study patterns of introgression (Hufford et al. 2013; Suarez-Gonzalez et al. 2016; Medugorac et al. 2017), to map genes involved in reproductive isolation (Corbett-Detig and Nielsen 2017) and phenotypic variation (Lindtke et al. 2013; vonHoldt et al. 2016), and to decipher past admixture processes (Brandvain et al. 2014; Liu et al. 2014). Although various LAI software are now available, they have been mainly tuned to human data in order to map disease-associated variants (Patterson et al. 2004; Seldin et al. 2011).

**Figure 1:**
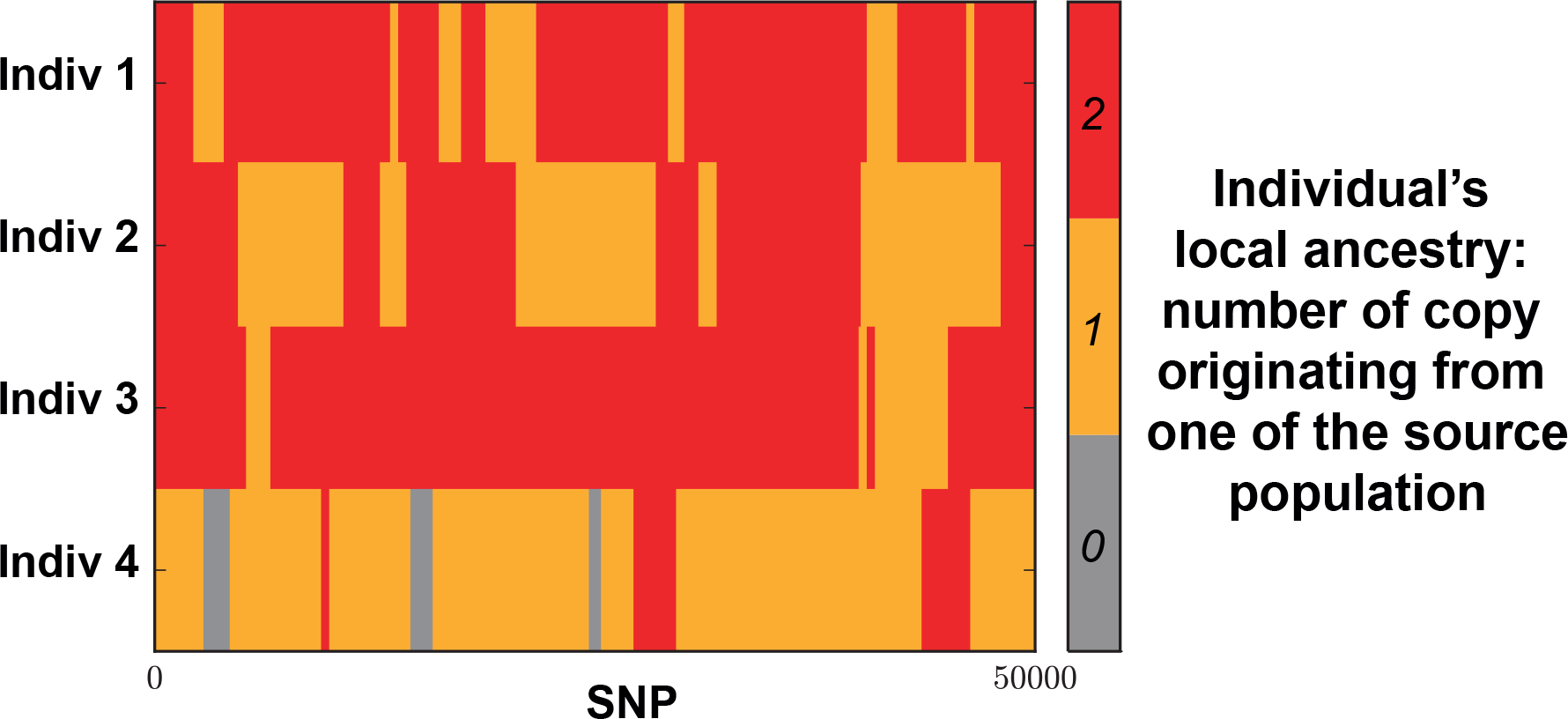
Example of local ancestry inference for 4 simulated Populus individuals resulting from admixture between 2 Populus species, which are *Populus trichocarpa* and *Populus balsamifera* (Suarez-Gonzalez et al. 2016). For an admixed individual, local ancestry at a given locus corresponds to the number of copies that has been inherited from the species *P. trichocarpa*. LAI software require haplotypes from putative source populations and process haplotypes or genotypes from admixed population to return local ancestry of admixed individuals. Details of the simulations are described in the Materials and Methods section.

In this paper, we introduce the software package Loter for Local Ancestry Inference, which does not require specifications of statistical or biological parameters in order to make LAI available for a wide range of species. Several software packages for LAI have been developed including HAPMIX, LAMP-LD, and RFMIX (Price et al. 2009; Baran et al. 2012; Maples et al. 2013). However, they require various parameters to be specified, which can hamper their practical use. HAPMIX requires specifications of several biological parameters that might be difficult to obtain such as genetic maps, recombination and mutation rate, average ancestry coefficients, and the average number of generations since admixture (Price et al. 2009). LAMP-LD requires a physical map and statistical parameters such as the number of hidden states in the hidden Markov Model and a window size where local ancestry is assumed to be constant (Baran et al. 2012). Default parameter values can be provided when using LAMP-LD. RFMIX requires statistical and biological parameters, which are a genetic map, a window size (in centimorgan) where local ancestry is assumed to be constant, and the average number of generations since admixture (Maples et al. 2013). Except for the genetic map, default parameters values can also be provided when using RFMIX (Maples et al. 2013). Other less-important differences between LAI software packages are also provided in Table 1. Except for LAMP-LD that uses statistical parameters only, RFMIX and HAPMIX require biological information such as genetic map, which can be difficult to provide for non-model species.

**Table 1:** Differences between several LAI software. The abbreviation param. stands for parameter and pop. for population. Other biological parameters required by HAPMIX are recombination and mutation rates.

|  | HAPMIX | LAMP-LD | RFMix | Loter |
| --- | --- | --- | --- | --- |
| Phasing of admixed individuals | No | No | Yes | Yes |
| Number of ancestral pop. | 2 | 2, 3, 5 | $\geq 2$ | $\geq 2$ |
| Genetic map required (in cM) | Yes | No (Physical position) | Yes | No (Ordered SNPs) |
| Limitation of the number of SNPS | No | 50,000 | No | No |
| Phasing error correction | No | Not required | Yes | Yes for 2 ancestral pop. |
| Parallel implementation | No | No | Yes | Yes |
| Admixture time required | Yes | No | Yes | No |
| Other biological param. required | Yes | No | No | No |

The software package Loter is based on the copying model introduced by Li and Stephens (2003) and already used in HAPMIX to perform LAI (Price et al. 2009). The copying model assumes that given a collection of “parental” haplotypes from putative source populations, haplotypes from admixed individuals are modeled as a mosaic of existing parental haplotypes (Figure 2). The main difference with HAPMIX is that Loter is not based on a probabilistic formulation that requires specifying several parameters. Instead, Loter is based on an optimization problem parametrized with a single regularization value λ that penalizes switches between parental haplotypes. Solutions of the optimization problem are found by using a dynamic programming algorithm, whose computational complexity is linear with respect to the number of markers and the number of individuals from the source populations. Inference of local ancestry is found by averaging results obtained for different values of the regularization parameter λ and various runs of the algorithm.

**Figure 2:**
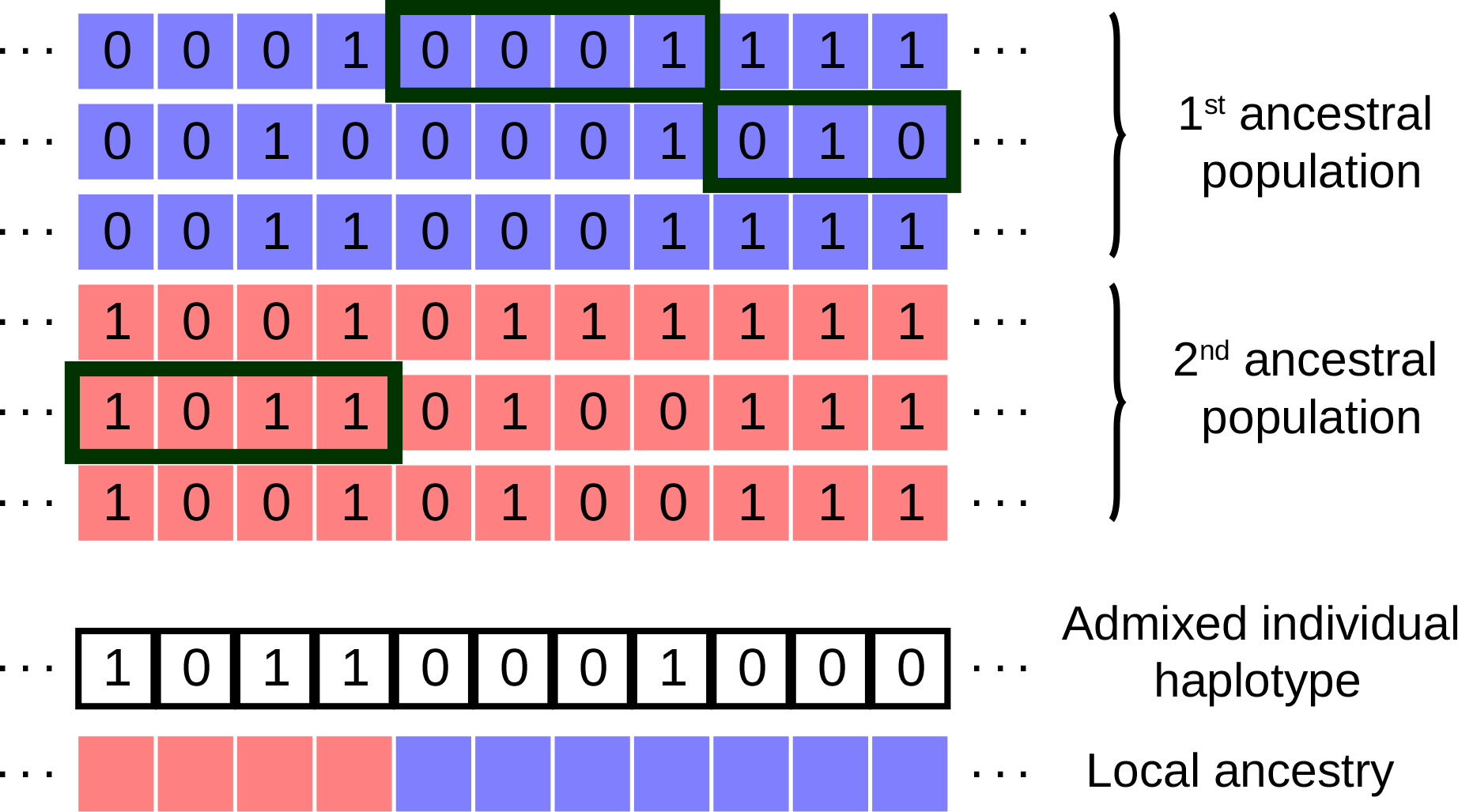
Graphical description of Local Ancestry Inference as implemented in the software Loter. Given a collection of parental haplotypes from the source populations depicted in blue and red, Loter assumes that an haplotype of an admixed individuals is modeled as a mosaic of existing parental haplotypes. In this example, the 1^st^ term of equation (1) (loss function) is equal to 1 because of a single mismatch between parental and admixed haplotype located at the next-to-last position, and the 2^nd^ term of equation (1) (regularization term) is equal to 2 λ because there are 2 switches between parental haplotypes. The displayed solution corresponds to the mathematical solution (*s*_1_, …, *s*_11_) = (5, 5, 5, 5, 1, 1, 1, 1, 2, 2, 2) where haplotypes are numbered from top to bottom, and *s_j_* = *k* if the admixed haplotype results from a copy of the *k*^th^ parental haplotype at the *j*^th^ SNP.

We compare Loter to HAPMIX, LAMP-LD, and RFMIX using diploid accuracy, which is an error measure analogous to imputation error for LAI (Sankararaman et al. 2008b). We consider an admixture experiment using human genotypes from HAPMAP 3 where we simulate genotypes resulting from admixture between Europeans (CEU) and Africans (YRI) (International HapMap 3 Consortium et al. 2010). We evaluate to what extent diploid accuracy is affected by the number of generations since admixture for the different approaches. We additionally consider a 3-way admixture scenario between Chinese (CHB), Europeans (CEU), and Africans (YRI) from HAPMAP 3. We repeat admixture simulations using genotypes from two Populus species in North America (Figure 1) (Suarez-Gonzalez et al. 2016). We additionally evaluate to what extent length of ancestry tracts are accurately inferred by the different LAI software packages. Length of ancestry tracts is a biological information that is used to date and reconstruct admixture events and that should consequently be accurately inferred for reliable demographic reconstruction (Gravel 2012; Ni et al. 2016; Corbett-Detig and Nielsen 2017; Xue et al. 2017). Finally, we apply Loter to admixed Populus genotypes, and we estimate admixture time using length of reconstructed ancestry tracts.

## New Approaches

We describe the optimization problem, which accounts that haplotypes from admixed individuals are described as a mosaic of haplotypes originating from the source populations (Figure 2). We assume that there are *n* individuals in the source populations resulting in 2*n* haplotypes denoted by (*H*_1_, …, *H*_2*n*_). The value (0 or 1) of the *i^th^* haplotype at the *j^th^* SNP is denoted by 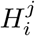. Haplotypes can be obtained from genotypes using computational phasing software such as fastPHASE or Beagle (Scheet and Stephens 2006; Browning and Browning 2007). A vector (*s*_1_, …, *s_p_*) describes the sequence of haplotype labels from which the haplotype *h* of an admixed individual can be approximated (Figure 2). For the *j^th^* SNP in the data set, *s_j_* = *k* if haplotype *h* results from a copy of haplotype *H_k_*. The optimization problem consists of minimizing the following cost function

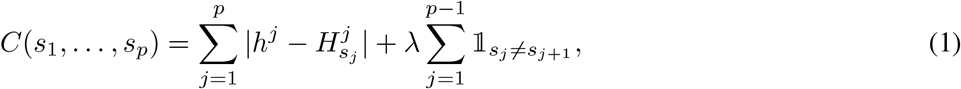

where (*s*_1_, …, *s_p_*) is in {1, · · ·, 2*n*}*^p^*. The first term in equation (1) is a loss function that sums over all possible loci of a {0, 1}-valued function equals to 1 if haplotype *h* is different from the copied haplotype and to 0 otherwise. The second term is a regularization function that is equal to the regularization parameter λ times the number of switches between parental haplotypes. The optimization problem of equation (1) can be visualized using the graph of Figure 3. A solution to minimize equation (1) can be found using dynamic programming and is provided in the Materials and Methods section. Once a solution has been provided about the sequence (*s*_1_, …, *s_p_*) of parental haplotypes, local ancestry values can be deduced automatically from this sequence because each parental haplotype belongs to one of the source populations (Figure 2). The formulation described in equation (1) is valid for *K* = 2 or more ancestral source populations.

**Figure 3:**
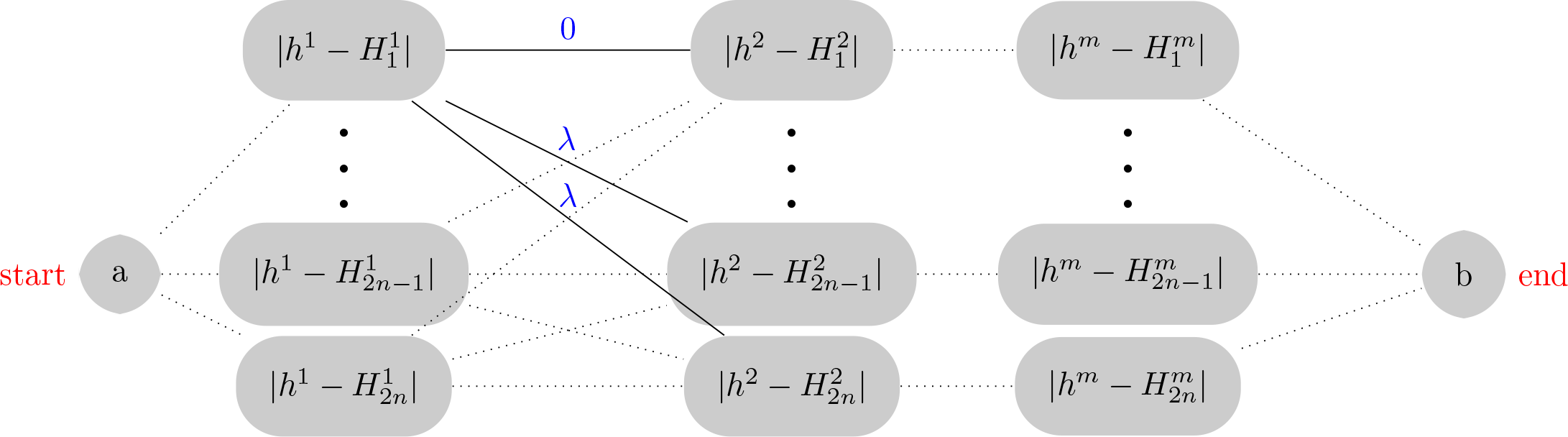
Graph that represents the optimization problem of equation (1). An optimal solution for (*s*_1_, …, *s_p_*) is found by finding the shortest path from node *a* to node *b*. We assume that there are *n* individuals in the source populations resulting in 2*n* haplotypes denoted by (*H*_1_, …, *H*_2*n*_). The value (0 or 1) of the *i^th^* haplotype at the *j^th^* SNP is denoted by 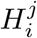. A vector (*s*_1_, …, *s_p_*) describes the sequence of haplotype labels from which the haplotype *h* of an admixed individual can be approximated. For the *j^th^* SNP in the data set, *s_j_* = *k* if haplotype *h* results from a copy of haplotype *H_k_*.

The optimization problem described in equation (1) involves a regularization parameter λ. Large values of λ strongly penalize switches between parental haplotypes such that solutions have long chunks of constant local ancestry. To avoid the difficult choice of λ, solutions for local ancestry are averaged by running the optimization method several times on different values of λ. Besides, to improve the stability of the solution, for each value of λ, we consider a bagging technique where 20 different solutions are found based on 20 different datasets generated using bootstrap with resampling (Breiman 1996). We consider λ = 1 + 0.5*i*, *i* = 1, …, 8, resulting in 160 = 8 *×* 20 different solutions and the final choice for local ancestry is obtained using a majority rule. When the most frequent vote has less that 75% of the votes, ancestry is imputed using local ancestry values of the closest SNPs with a preference for the SNP on the left in case of ambiguity. Finally, an additional smoothing procedure is considered in order to account for switch errors when phasing admixed individuals (see Materials and Methods).

## Results

### Simulated human admixed individuals

We consider simulated admixed individuals resulting from admixture between Africans (YRI population in HAPMAP 3) and Europeans (CEU population in HAPMAP 3). Accuracy obtained with several LAI software packages varies depending on time since admixture occurs (Figure 4). For recent admixture where admixture occurred 5 generations ago, LAMP-LD and RFMix obtain the best result with median diploid accuracies of 99.6% and 99.8% whereas median diploid accuracy of Loter is equal to 99.3%. For ancient admixture where admixture occurred 500 generations ago, Loter obtains the largest median diploid accuracy of 86.7% followed by LAMP-LD with a diploid accuracy of 80.6% and RFMix with a diploid accuracy of 72.0%. For the smallest admixture times (*µ* ≤ 20 generations), diploid accuracy obtained with the three approaches is larger than 95% and RFMix and LAMP-LD outperform LOTER. By contrast, for the largest admixture times (*µ* ≥ 150 generations), Loter is the most accurate software for LAI (Figure 4).

**Figure 4:**
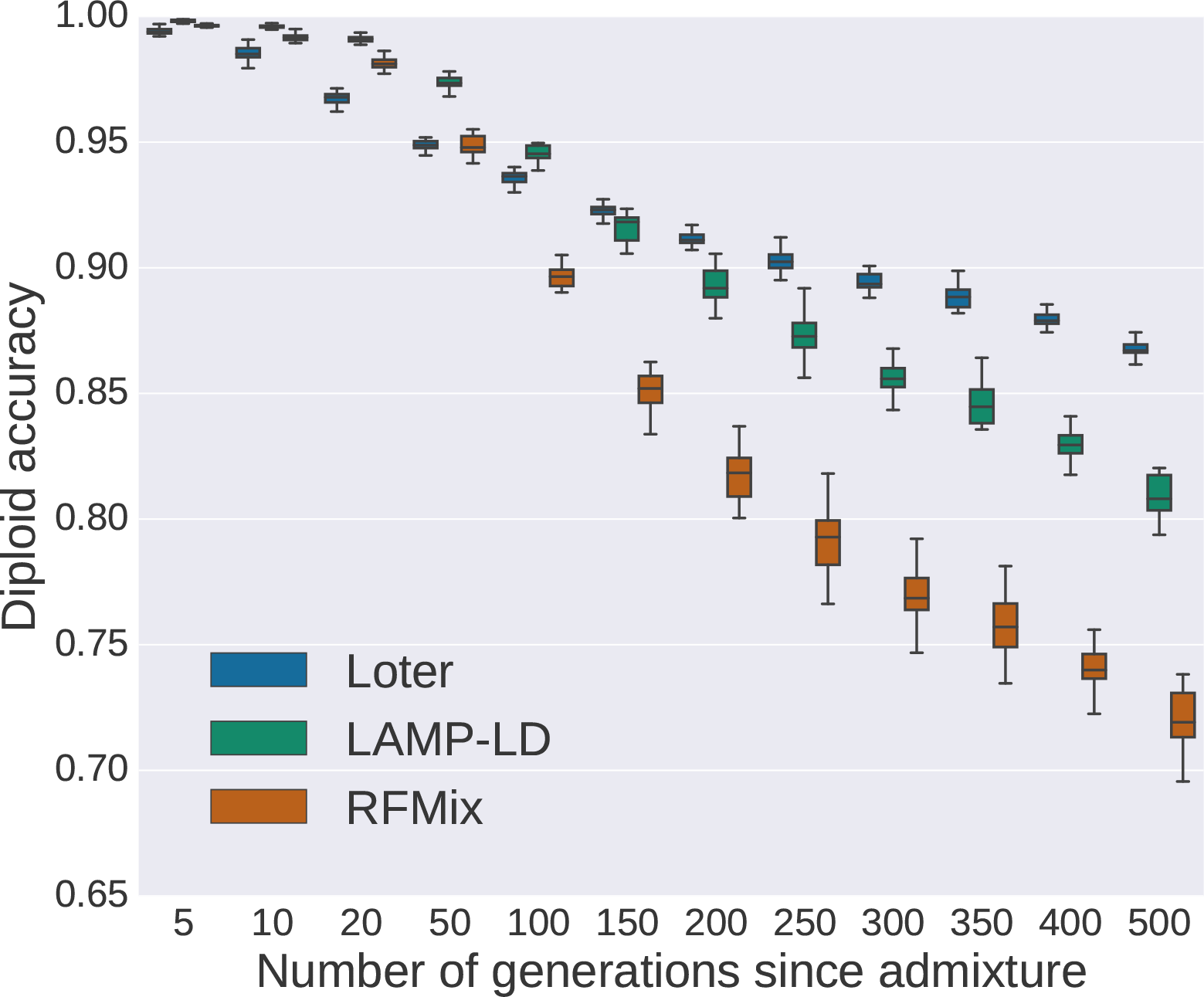
Diploid accuracy obtained with LAMP-LD, Loter, and RFMix for simulated admixed human individuals as a function of the time since admixture occurred. Admixed individuals are simulated by constructing their genomes from a mosaic of true African (YRI) and European (CEU) haplotypes (International HapMap 3 Consortium et al. 2010). For performing simulations, true haplotypes are obtained using trio information. For local ancestry inference, haplotypes are obtained with Beagle using individuals that are not used for simulating admixed individuals. For each value of the number of generations since admixture, 20 sets of 48 admixed individuals are generated. Boxplots show the distribution of the 20 values for the mean diploid accuracy.

**Figure 5:**
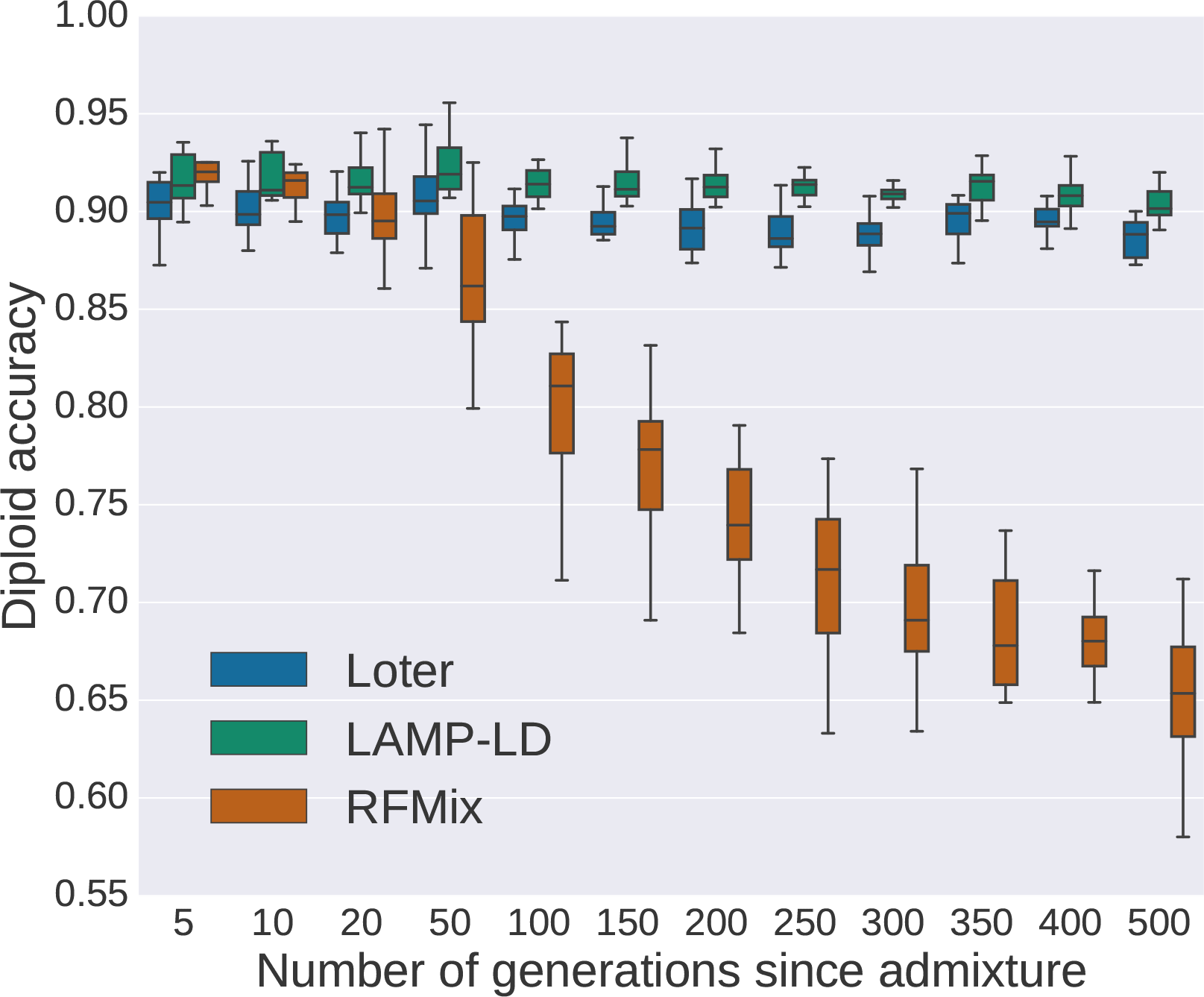
Diploid accuracy obtained with LAMP-LD, Loter, and RFMix for simulated admixed Populus individuals as a function of the time since admixture occurred. Admixed individuals are simulated by constructing their genomes from a mosaic of *Populus trichocarpa* and *Populus balsamifera* individuals. Individuals are phased using Beagle and two different sets of individuals are used for performing simulations and inference. For each value of the number of generations since admixture, 20 sets of 20 admixed individuals are generated. Boxplots show the distribution of the 20 values for the mean diploid accuracy.

To evaluate these LAI software packages in a different context, we consider an additional setting where haplotypes of the reference panel are not drawn from the source populations but from populations that have diverged from the source populations. For individuals resulting from admixture between Yoruban (YRI) and Northern-European ancestry (CEU), reference panels are the Luhya sample from Kenya (LWK) and the Toscani sample from Italy (TSI). Compared to Loter, we again find that RFMix and LAMP-LD provide more accurate reconstructions of local ancestry for the recent admixture times of *µ* = 5, 10, 20 generations but less accurate reconstructions for more ancient admixture events (Supplementary Figure SI1). Loter is more accurate then RFMix when *µ* ≥ 50 generations; e.g. Loter accuracy is of 92% for *µ* = 100 generations whereas RFMix accuracy is equal to 87%. The loss of accuracy of LAMP-LD for ancient admixture times is less pronounced but Loter still provides more accurate reconstructions than LAMP-LD when *µ* ≥ 100 generations.

We evaluate the benefit of bagging and of averaging local ancestry values obtained with different values of the regularization parameter, which are implemented by default in Loter. First, diploid accuracy depends on the choice of the regularization parameter (Supplementary Figure SI2). As expected, smallest values of provide the best result for ancient admixture event; λ = 2 is optimal when *µ* = 400 and *µ* = 500 generations, and λ = 5 is optimal otherwise. Second, for all values of the admixture times, averaging local ancestry values obtained with different values of the regularization parameter λ instead of considering a single value improves inference (Supplementary Figure SI2). Last, averaging over bootstrap replicates (bagging) further improves local ancestry inference (Supplementary Figure SI2). Additionally, we vary the number of haplotypes in the reference panels to investigate to what extent Loter is sensitive to the reference panels. When using the TSI and LWK reference samples, the diploid accuracy varies substantially from 41% with a single haplotype in each population to 98% when using 100 haplotypes in each population.

We additionally evaluate the diploid accuracy of HAPMIX, which is another LAI software package based on the copying model. Compared to the three other LAI software packages, its diploid accuracy is the smallest with values of diploid accuracy ranging from 42% to 57% (Supplementary Figure SI3). The fact that haplotypes used for LAI are not exact but can be computationally phased using Beagle causes the severe reduction of diploid accuracy obtained with HAPMIX (Supplementary Figure SI4). When considering haplotypes reconstructed with Beagle instead of true haplotypes, diploid accuracy is reduced by 32% on average.

Additionally, we compare diploid accuracy of Loter, RFMix and LAMP-LD on data simulated under a 3-way admixture model (Supplementary Figure SI5). As for 2-way admixture model, we find that RFMix and LAMP-LD have larger diploid accuracies than Loter for recent admixture and smaller ones for ancient admixture. When admixture took place 5 generations ago, the diploid accuracy of RFMix is of 99.8%, the accuracy of LAMP-LD is of 99.8%, and the accuracy of Loter is of 99.5%. By contrast, when admixture took place 500 generations ago, the diploid accuracy of RFMix is of 84.6%, the accuracy of LAMP-LD is of 89.1%, and the accuracy of Loter is of 92.3%.

### Simulated Populus admixed individuals

We simulate individuals that result from admixture between *Populus trichocarpa*, which is adapted to relatively humid, moist, and mild conditions west of the Rocky Mountains and *Populus balsamifera*, which is a boreal species (Suarez-Gonzalez et al. 2016). As for human data, we compare Loter, RFMix, and LAMP-LD using diploid accuracy as a criterion for comparison. Again, the diploid accuracy of RFMix decreases with increasing admixture time. It ranges from a diploid accuracy of 92.0% when admixture occurs 5 generations ago to a diploid accuracy of 65.3% when admixture occurs 500 generations. By contrast to the simulations of human data, we find that the diploid accuracy of Loter and of LAMP-LD does not change with admixture time. For LAMP-LD, diploid accuracies ranges from 91.3% to 90.1% and for Loter it ranges from 89.9% to 89.0% when admixture increases from 5 to 500 generations.

### Length of ancestry tracts

For simulated admixed Populus individuals, we additionally compare the length of *P. balsamifera* reconstructed ancestry tracts to the length of true ancestry tracts (Figure 6). When admixture took place 10 generations ago, RFMIX provides a distribution of ancestry tracts that is closer to the true distribution. For Populus simulations, the true median length of ancestry tract is of 4.26 cM and RFMIX finds 5.20 cM whereas LAMP-LD and Loter find median lengths of 0.05 and 2.31 cM respectively. Both LAMP-LD and Loter return spurious chunks of local ancestry of small lengths and that contribute to reduce the mean length of local ancestry (Supplementary Figure SI6). Additionally, several long blocks of ancestry chunks are cut into smaller pieces when using Loter or LAMP-LD (Supplementary Figure SI6). When admixture took place 10 generations ago in the human simulations, both Loter and RFMIX provide the most accurate results; the true median length of ancestry tract is of 9.03 cM and RFMIX, Loter, and LAMP-LD reconstruct ancestry tracts of median length 9.99 cM, 12.20 cM and 9.02 cM, respectively.

**Figure 6:**
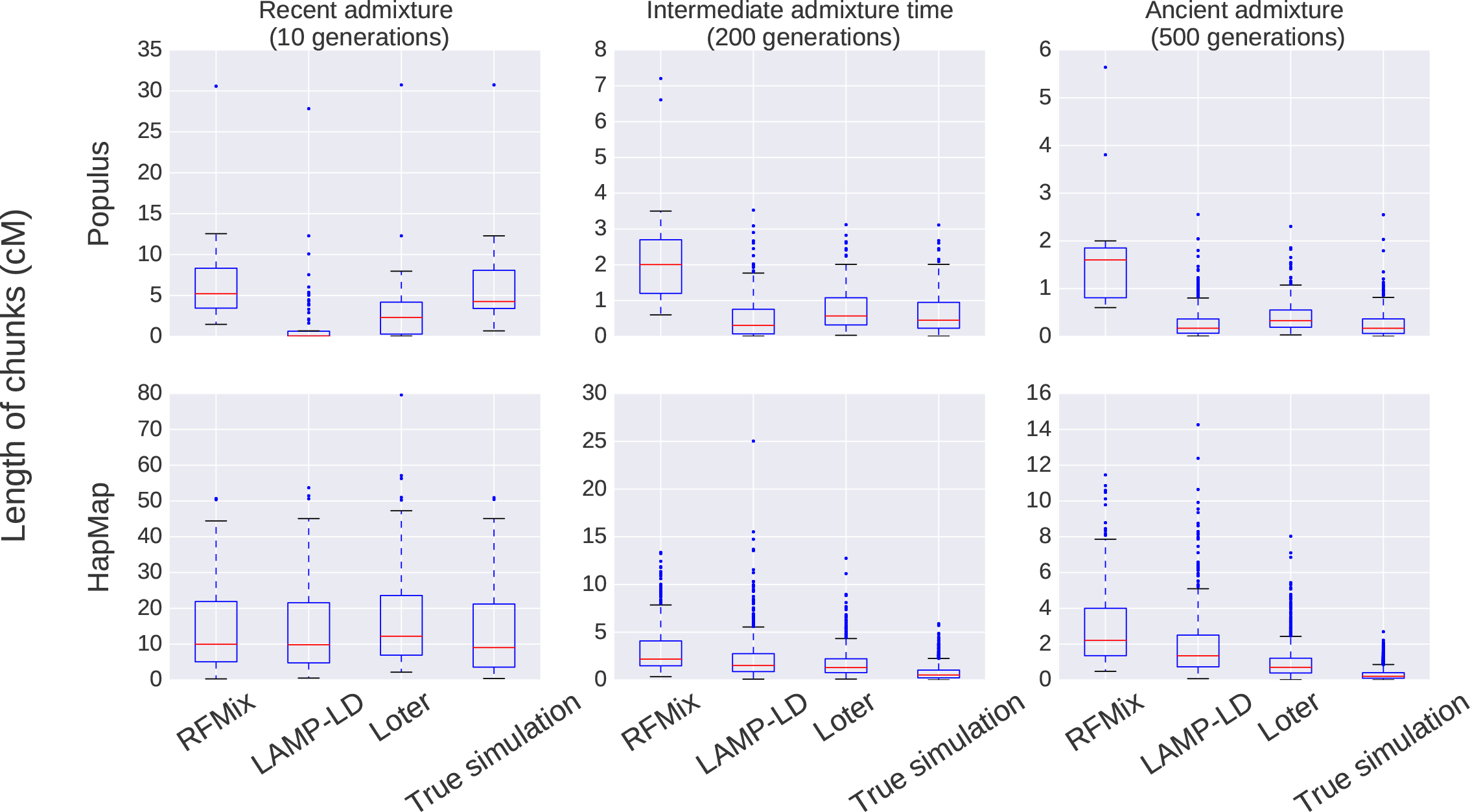
Distribution of the length of ancestry chunks for simulated data. For Populus data, we consider the first 500,000 SNPs of chromosome 6 and for human data, we consider the first 50,000 SNPs of chromosome 1. When considering Populus data, we run 10 times LAMP-LD on non-overlapping sets of SNPs in order to avoid the limitation of 50,000 SNPs of LAMP-LD.

When admixture took place 200 or of 500 generations, length of ancestry chunks are more accurately reconstructed with Loter and with LAMP-LD than with RFMIX (Figure 6). For both human and Populus simulated data, RFMix, by contrast to Loter and LAMP-LD, reconstructs ancestry tracts that are too long compared to true ancestry tracts when admixture is larger or equal than 200 generations. When admixture took place 200 generations ago, true median ancestry tracts is of 1 cM or less whereas RFMix reconstructs tracts of 2 cM or more. For Populus simulations, the true median length of ancestry tract is of 0.45 cM when admixture took place 200 generations ago, and RFMIX, Loter, and LAMP-LD find respectively 2.00 cM, 0.56 cM and 0.30 cM (Supplementary Figure SI6). When admixture took place 500 generations ago, the true median length of ancestry tract is of 0.17 cM, and RFMIX, Loter, and LAMP-LD find respectively 1.60 cM, 0.33 cM and 0.17 cM.

### Application of Loter to admixed Populus individuals

We applied Loter to 36 individuals that are admixed between *P. balsamifera* and *P. trichocarpa*. When averaging local ancestry coefficients, we find that admixed individuals have on average 87% of *P. trichocarpa* ancestry and 13% of *P. balsamifera* ancestry. We find that the median length of *P. balsamifera* ancestry tracts is equal to 0.76 cM and the first and third quartiles are equal to 0.25 cM and 1.47 cM. We also perform simulations of admixed individuals based on true genotypes from *P. balsamifera* and *trichocarpa* individuals. When admixture time varies from 10 to 500 generations; median reconstructed *P. balsamifera* ancestry tracts vary from 2.3 cM to 0.32 cM, the first quartile varies from 0.28 cM to 0.19 cM, and the third quartile varies from 4.18 cM to 0.55 cM (Figure 7). Of the six admixture times we considered (*µ* ∈ {10, 50, 100, 200, 300, 500}) in the simulations, we find that *µ* = 100 generations provide the most similar distribution of *P. balsamifera* ancestry tracts; when *µ* = 100 generations, the three quartiles of *P. balsamifera* ancestry tracts are equal to 0.21 cM, 0.78 cM, and 1.29 cM.

**Figure 7:**
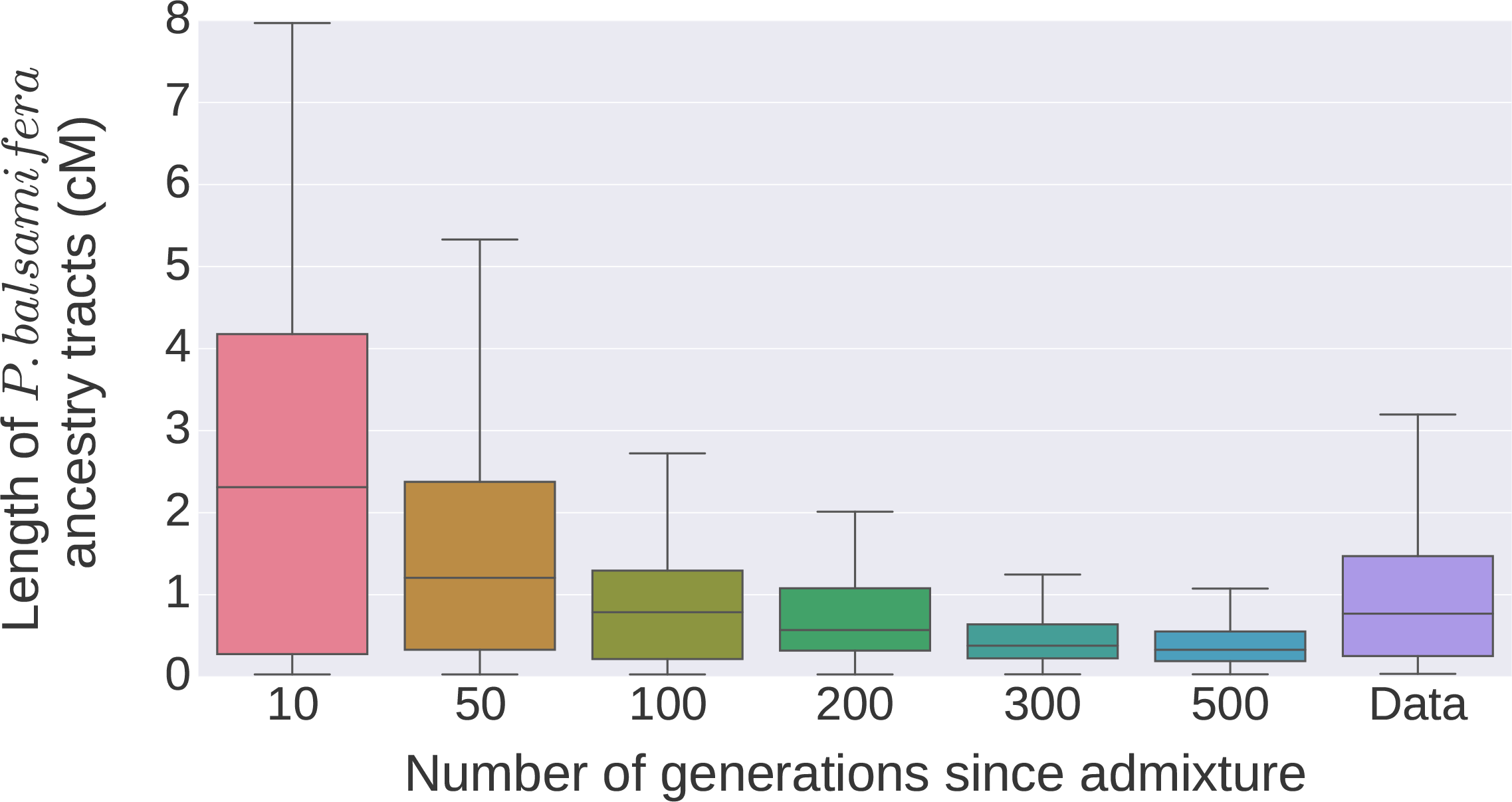
Distribution of the length of *P. balsamifera* ancestry tracts. The data consist of genotypes of admixed individuals between *P. balsamifera* and *P. trichocarpa*. For the simulations, we replicate the same pipeline as for local inference with real data, which consist of using Beagle to phase genotypes and Loter to reconstruct ancestry tracts.

## Discussion

As dense genotype or sequencing data become more affordable, local ancestry inference provides an opportunity for admixture mapping and for deciphering admixture processes as well. We have introduced the software package Loter in order to make local ancestry available for a wide range of species for which biological parameters such as admixture times or recombination rates are not available. The regularization parameter λ, which controls smoothing, depends in a complicated manner on several biological and statistical parameters including mutation and recombination rates. To avoid the difficult choice of the regularization parameter, Loter implements an averaging procedure where we average solutions for different values of the regularization parameter. Thanks to an appropriate model averaging strategy, Loter does not require parameter tuning which makes it easy to use.

Loter requires phased haplotypic data for both the target as well as the reference panels. Usually haplotype phases are found using computational phasing programs such as Beagle or shapeIT that may have different phasing accuracies (see Fig. S. 24 of Malinsky et al. 2017). To investigate effects of phasing on ancestry reconstruction, we have conducted an additional experiment using the CEU-YRI admixture simulations (admixture time of 5 generations). We have modified haplotype phases of target and admixed samples until there is a 30% switch error rate with the true haplotypic phase. We found that diploid accuracy is the same (97%) whether or not true haplotype phases are used, which shows that Loter is able to account for phasing errors.

Loter is based on a mathematical formulation, which involves minimizing an objective function using dynamic programming. The Viterbi algorithm, which is a particular instance of dynamic programming, has already been used to reconstruct local ancestry in Hidden Markov models (HMM) (Baran et al. 2012; Sankararaman et al. 2008a). Compared to other HMM formulations of local ancestries, Loter formulation is more parsimonious because it involves a single regularization parameter (Scheet and Stephens 2006; Baran et al. 2012). In addition to local ancestry reconstruction, dynamic programming has also been used to partition chromosomes into haplotype blocks, which correspond to discrete blocks of limited haplotype diversity (Kimmel et al. 2003; Zhang et al. 2002).

We compared Loter to the local ancestry software packages HAPMIX, RFMIX, and LAMP-LD. We found that the diploid accuracy obtained with HAPMIX is reduced by 32% in average when haplotypes are not known perfectly using trio-phasing but only reconstructed using phasing software such as Beagle. By contrast, RFMIX, LAMP-LD, and Loter are robust to imperfect haplotype reconstruction, which explains why only RFMIX, LAMP-LD, and Loter were further considered in software comparisons. When admixture took place 5 generations ago, RFMIX and LAMP-LD provide the largest diploid accuracies but all three software were found to provide an accurate reconstruction of local ancestry with diploid accuracies always larger than 99% for the simulations of Afro-American admixed haplotypes and larger than 89.9% for the simulations of Populus individuals. Compared to RFMIX and LAMP-LD, results obtained with Loter are more robust with respect to the time since admixture occurred. For simulated human data, the diploid accuracy of Loter decreases to 87% for the most ancient admixture times of 500 generations whereas it decreases to 72% and 81% when using RFMIX and LAMP-LD. For Populus data, diploid accuracy does not depend on admixture times when using Loter and LAMP-LD whereas it is severely impacted when using RFMIX. The fact that diploid accuracy is not impacted by the considered range of admixture times is encouraging and suggests that local ancestry inference is possible for admixture that occurred hundreds of generation ago when SNP density is large enough, as for Populus data; the mean distance between two SNPs being of 9.8.10^*−*6^ cM for Populus data and of 2.7.10^*−*3^ cM for the human data. A large enough genetic differentiation between parental populations is another explanation about why LAI can be accurate for Populus even for ancient admixture events; mean *F_ST_* between *Populus trichocarpa* and *Populus trichocarpa* is equal to 0.22 (A. Geraldes, personal communication) whereas it is only of 0.14 when comparing CEU and YRI populations (Bhatia et al. 2013).

Diploid accuracy may be impacted by admixture time and explanations may be related to the smoothing procedure of the different software. For instance, RFMIX has been tuned to genotypes resulting from recent admixture that occurred 10 generations ago or less such as admixture between African and Europeans (Gravel 2012; Bryc et al. 2015). When using default parameters of RFMIX for data resulting from ancient admixture events, reconstructed ancestry tracts are inadequately long (Figure 6 and Supplementary Figure SI6) and over-smoothing can affect diploid accuracy. Even when providing true admixture time to RFMIX, diploid accuracy of RFMIX remains more impacted by admixture time possibly because of the default of 0.2 cM for window size (Supplementary Figure SI7). When using HAPMIX, a choice of window lengths or of the time since admixture should also be provided. However, choice of admixture time can be very difficult to estimate and impacts biological results. For instance, lengths of ancestry tracts found with HAPMIX depend on the input value for admixture time (Patin et al. 2014). The model averaging procedure implemented in Loter has the advantage to avoid putting a strong prior on a particular length of ancestry tract. In addition, model averaging improves parameter inference, which has already been observed when phasing genotypes using a statistical model of Linkage Disequilibrium (Scheet and Stephens 2006).

The simulation results show that accuracy obtained with Loter and LAMP-LD is less sensitive to admixture times compared to the accuracy obtained with RFMix. LAMP-LD is more accurate that Loter for recent admixture times when using human data and for all values of admixture times when considering the Populus data. However, LAMP-LD has limitations for large-scale NGS data that contains a large number of molecular markers. Its current implementation has a memory allocation limit of 50,000 SNPs and the software can be computer intensive. To perform local ancestry inference for 500,000 SNPs and 20 admixed Populus individuals, we compare the running time of the three LAI software using 2 multi-core Intel Xeon E5645 2.40 GhZ processors (6 cores each); the running time is of 28 minutes using RFMIX, of 58 minutes with LAMP-LD and of 6 minutes using Loter. This small running time shows that Loter can handle massive genomic dataset using a desktop or laptop computer. However, although there are differences between LAI software packages, using different LAI software packages can be a wise strategy to provide enhanced evidence for an association or a selection signal (Zhou et al. 2016). We expect that providing a parameter-free and rapid software for local ancestry inference will make more accessible genomic studies about admixture processes.

## Materials and Methods

### Dynamic Programming

The optimization problem of equation (1) is solved using dynamic programming. The solution of the problem with *p* SNPs can be derived from the solution with (*p* − 1) SNPs. Two configurations are possible. Either the admixed haplotype copies from the same haplotype at the (*p* − 1)^th^ and *p*^th^ SNP and

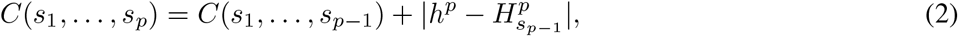

or it uses different template haplotypes and

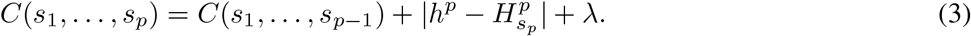

The optimal solution is then found by computing the shortest path on a graph displayed in Supplementary Figure 3. To find the shortest path, dynamic programming computes at each node a quantity *Q*(*i, j*) that corresponds to the optimal solution for the first *j* SNPs and when the template haplotype at SNP *j* is the *i^th^* haplotype *s_j_* = *i*. The or it uses different template haplotypes and quantity *Q*(*i, j*) is updated as followed

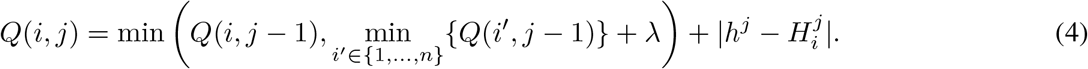

Because we store the value of min_*i*′ ∈{1,…,*n*}_{*Q*(*i′, j* − 1)} at locus *j* − 1, the value of *Q*(*i, j*) can be computed as a minimum between 2 values. For each admixed haplotype, the complexity of this algorithm is therefore *𝒪*(*n × p*) where *n* is the number of individuals in the ancestral populations and *p* is the number of SNPs. The path (*s*_1_, …, *s_p_*) is then converted to an haploid ancestry sequence *a* = (*a*_1_, …, *a_p_*) ∈ (1, …, *K*)^*p*^ where *a_j_* is the population of origin of the 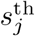haplotype.

### Accounting for phase errors of admixed genotypes

Reconstructed haplotypes from an admixed population may contain switch errors (Pasşaniuc et al. 2009; Browning and Browning 2011). As considered in RFMix, it is possible to redistribute ancestry chunks among the two haplotypes from the same individual in order to correct for switch errors. For now, the software Loter accounts for phase error when there are 2 ancestral populations only. Once local ancestry values for each of the 2 haplotypes have been found after solving equation (1), we compute the sum of the 2 haplotypic local ancestries resulting in diploid local ancestry values *d* = (*d*^1^, …, *d^p^*) ∈ {0, 1, 2}^*p*^ as returned by the software HAPMIX. Local ancestry values are then reconstructed using an internal *ancestry phasing* algorithm. The phasing procedure considers that the 2 haplotypic local ancestry sequences are a mosaic of two possible ancestry sequences *A*_1_ = (0, …, 0) and *A*_2_ = (1, …, 1) corresponding to the two possible ancestral populations. Two vectors (*s*_1_, …, *s_p_*) ∈ {0, 1}^*p*^ and (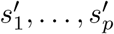) ∈ {0, 1}^*p*^ describe the sequence of labels, two vectors *a* ∈ {0, 1}^*p*^ and *a*′ ∈ {0, 1}^*p*^ are the haploid local ancestry values, and Θ is a compound parameter equal to (*s*_1_, …, *s*_*p*_, 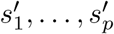, *a, a*′). The ancestry phasing algorithm consists of minimizing the following cost function

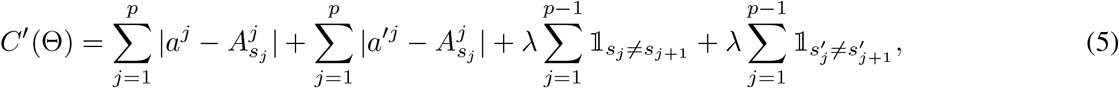

subject to the constraint that the sum of the haploid local ancestries *a* and *a′* is equal to the diploid local ancestry *d*. The solution for Θ is found using dynamic programming. For each admixed 𝒪(*p*). Once a solution has been found, haplotypic local ancestry, which has been corrected for phasing errors, consists of the two sequences (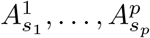 and 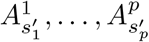).

### Parameter settings for local ancestry software

For RFMix, LAMP-LD, and HAPMIX, we use the default parameter settings.

### Admixture between Populus species

We simulate admixed individuals by constructing their genomes from a mosaic of real *P. balsamifera* and *P. trichocarpa* individuals (Suarez-Gonzalez et al. 2016). We consider a probabilistic model that has already been used to simulate admixed individuals and to evaluate the performances of HAPMIX (Price et al. 2009). Simulations are based on 20 haplotypes (first 50,000 SNPs) from chromosome 6 from the species *P. balsamifera* (balsam poplar) and of 20 haplotypes from the species *P. trichocarpa* (black cottonwood) (Suarez-Gonzalez et al. 2016). Haplotypes were obtained from genotypes using Beagle (Browning and Browning 2007). The *P. trichocarpa* ancestry *ɑ_i_* of a simulated admixed individual is drawn randomly according to a Beta distribution of mean 0.8 and of variance 0.1. At the first marker, the haplotype of an admixed individual *i* is assumed to originate from *P. trichocarpa* with a probability *ɑ_i_* and from *P. balsamifera* otherwise. For each simulated haplotype, we associate one *P. balsamifera* haplotype and one *P. trichocarpa* haplotype. For a given admixed individual, haplotypes are exclusively copied from these two source haplotypes that are chosen at random. The length (measured in Morgans) of an ancestry chunk is drawn according to an exponential distribution of rate *µ* generations. In the simulations, we consider values for *µ* ranging from 5 to 500 generations. The species origin of the new ancestry tract is again determined using the (*ɑ_i_*, 1 *− ɑ_i_*) admixture coefficients and the copying process for haplotype is repeated as before. To reconstruct local ancestry of simulated admixed individuals, we consider 30 haplotypes from *P. balsamifera* and 30 haplotypes from *P. trichocarpa* that were not used when simulating admixed individuals. Haplotypes were again phased using the software Beagle. To evaluate diploid accuracy for a given value of *µ*, we consider 20 sets of simulations consisting of 20 admixed haplotypes each.

### Admixture between human populations

When simulating admixed individuals between Yorubans (YRI) and Europeans (CEU) from HAPMAP 3, we consider the copying process mentioned before. For simulations, we consider true haplotypes based on trio phasing. For inference, we consider haplotypes reconstructed using Beagle based on genotypes that are not used for simulations. A total of 48 Yoruban haplotypes and of 48 European haplotypes are considered to simulate 48 Afro-American haplotypes. We consider 40 European haplotypes and 40 African haplotypes, which were obtained with Beagle, to perform local ancestry inference. To evaluate diploid accuracy for a given value of *µ*, we consider 20 sets of simulations consisting of 48 admixed haplotypes each. When considering reference panels different from the source populations, we consider 100 haplotypes from the HAPMAP 3 Luhya sample and 176 samples from the HAPMAP 3 TSI samples. Local ancestry reconstruction was based on true haplotypes from the LWK and TSI sample.

We consider an additional set of simulations where 3 populations admixed. Admixture is assumed to occur between Chinese (CHB), Europeans (CEU) and Africans (YRI). The exponential distribution for European chunks is of rate *µ* ∈ {5, 100, 200, 500} generations and the exponential distribution for African and Asian chunks is equal to *µ/*2 generations. By contrast to the 2-way admixture models, we use trio-phased haplotypes for inference and not haplotypes reconstructed with Beagle.

### Admixed Populus individuals

We consider 36 individuals that are admixed between *P. balsamifera* and *P. trichocarpa* (Suarez-Gonzalez et al. 2016) and use Beagle to phase them. We use the first 500, 000 SNPs of chromosome 6 to reconstruct ancestry tracts. We simulate 16 admixed individuals based on 20 genotypes from *P. balsamifera* and *P. trichocarpa* species. Instead of considering true ancestry ancestry tracts, we rather replicate the same pipeline as for real data such as the bias of ancestry tract reconstruction should be same for data and simulations. We phase simulated individuals using Beagle and reconstruct ancestry tracts using Loter.

### Software

The Loter software package and its source code are available at https://github.com/bcm-uga/Loter.

### Data

Human HapMap 3 data are available online at http://www.sanger.ac.uk/resources/downloads/human/hapmap3.html. Populus data are available online at http://datadryad.org/resource/doi:10.5061/dryad.0817m.

## Acknowledgments

Authors acknowledge Grenoble Alpes Data Institute, supported by the French National Research Agency under the “Investissements d’avenir” program (ANR-15-IDEX-02), the LabEx PERSYVAL-Lab (ANR-11-LABX-0025-01) and the MACARON project (ANR-14-CE23-0003). JM is also supported by a grant from the European Research Council (SOLARIS project, number 714381).

---

**Figure SI1:**
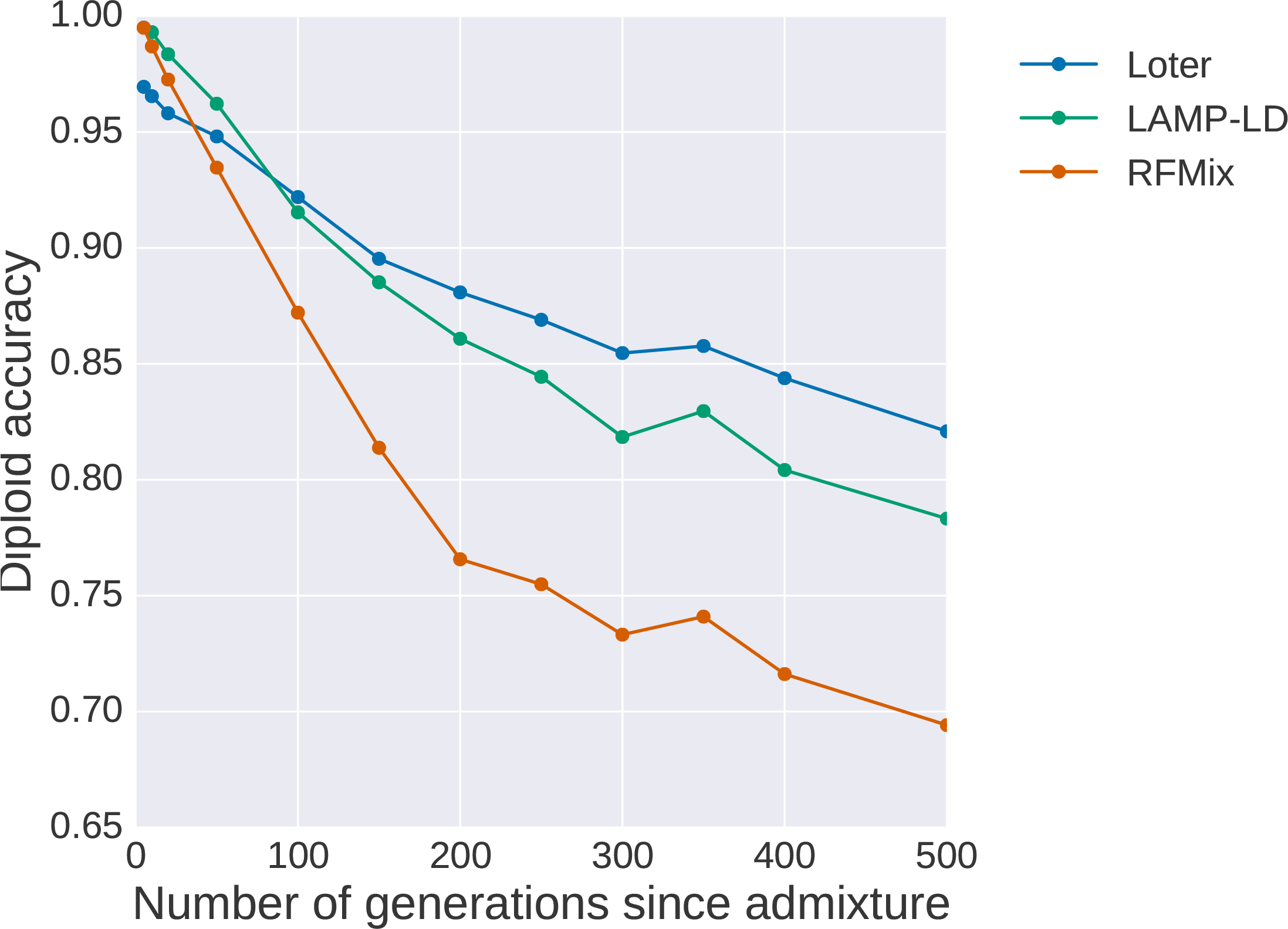
Diploid accuracy obtained with LAMP-LD, Loter, and RFMix when haplotypes of the reference panel are not drawn from the source populations but from populations that have diverged from the source populations. Admixed individuals are simulated by constructing their genomes from a mosaic of true African (YRI) and European (CEU) haplotypes and reference panels are the Luhya sample from Kenya (LWK) and the Toscani sample from Italy (TSI) (International HapMap 3 Consortium et al. 2010). For performing simulations, true haplotypes are obtained using trio information. For local ancestry inference, haplotypes are obtained with Beagle using individuals that are not used for simulating admixed individuals. For each value of the number of generations since admixture, 1 set of 48 admixed individuals are generated.

**Figure SI2:**
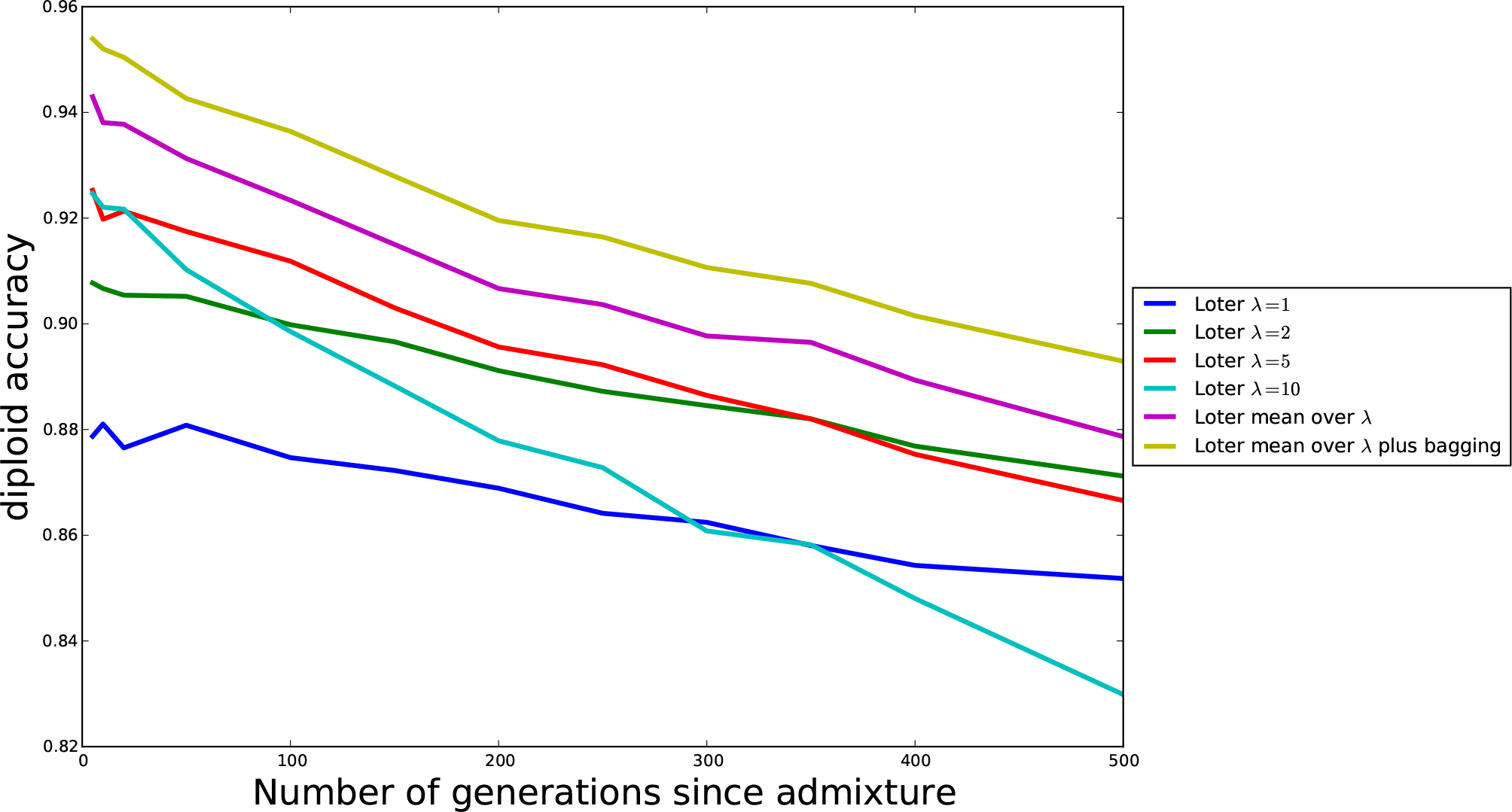
Diploid accuracy obtained with Loter is improved when bagging and when averaging over multiple values of the regularization parameter. Admixed individuals were simulated by constructing their genomes from a mosaic of true African (YRI) and European (CEU) haplotypes (International HapMap 3 Consortium et al. 2010) (Figure 4). Diploid accuracies are evaluated for twelve different values of the admixture time corresponding to 5, 10, 20, 50, 100, 150, 200, 250, 300, 350, 400 and 500 generations.

**Figure SI3:**
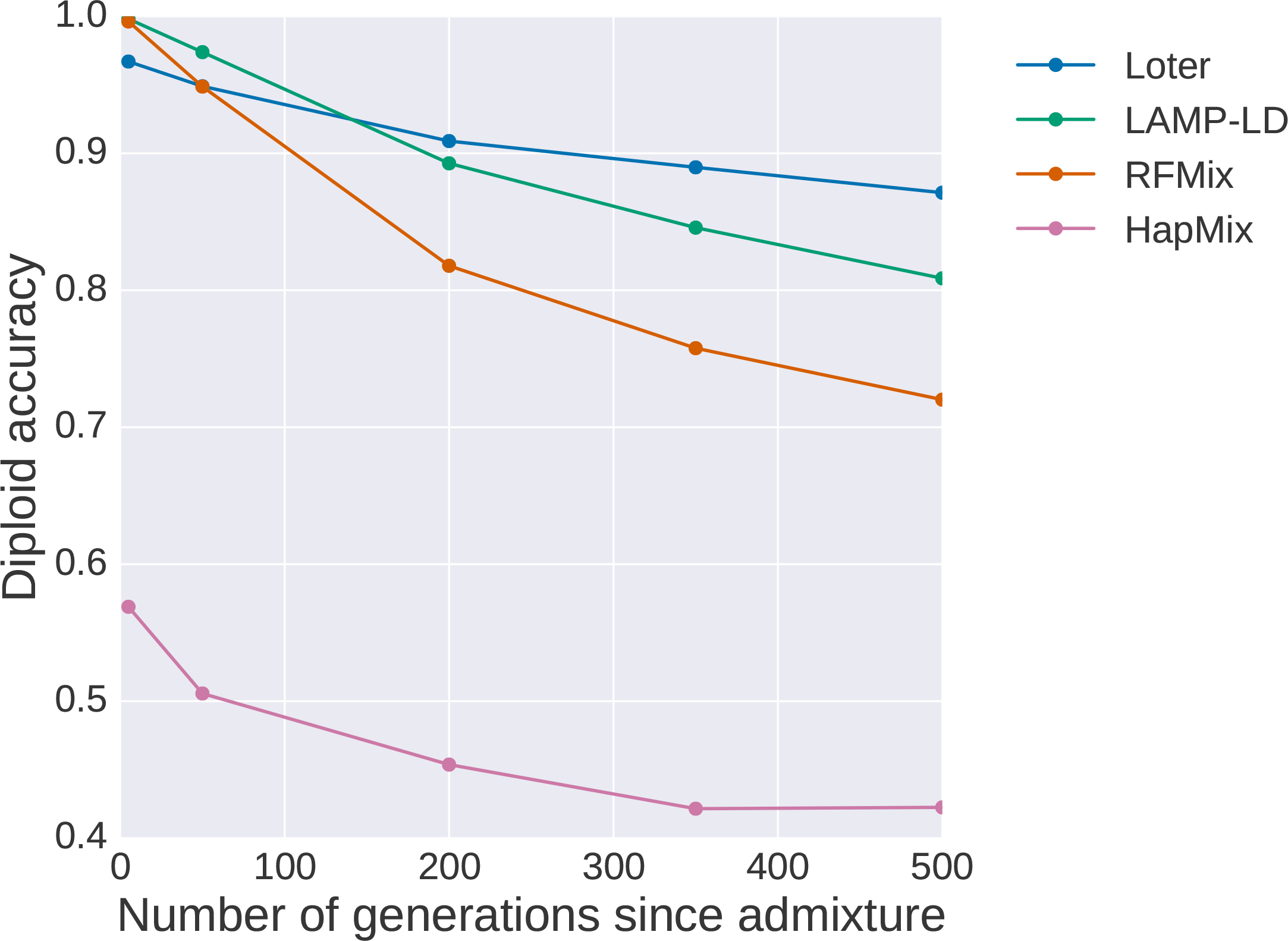
Diploid accuracy obtained with LAMP-LD, Loter, and RFMix for simulated human individuals as a function of the time since admixture occured. Admixed individuals are simulated by constructing their genomes from a mosaic of true African (YRI) and European (CEU) haplotypes (International HapMap 3 Consortium et al. 2010) (Figure 4). For each admixture time, HAPMIX is evaluated using a single simulation of 48 admixed individuals. Other software, which run faster, are evaluated based on the mean diploid accuracy obtained with 20 simulated sets of 48 admixed individuals.

**Figure SI4:**
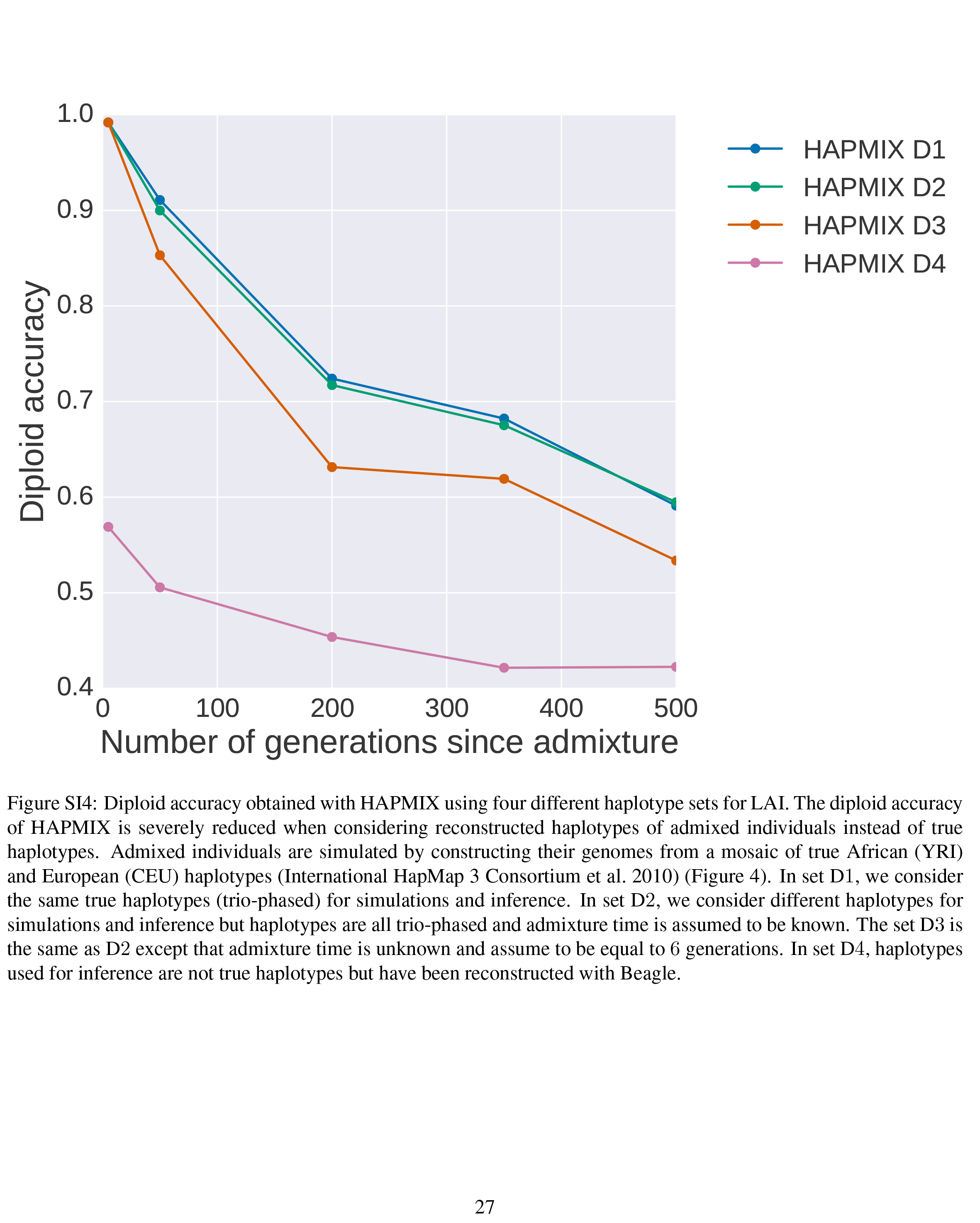
Diploid accuracy obtained with HAPMIX using four different haplotype sets for LAI. The diploid accuracy of HAPMIX is severely reduced when considering reconstructed haplotypes of admixed individuals instead of true haplotypes. Admixed individuals are simulated by constructing their genomes from a mosaic of true African (YRI) and European (CEU) haplotypes (International HapMap 3 Consortium et al. 2010) (Figure 4). In set D1, we consider the same true haplotypes (trio-phased) for simulations and inference. In set D2, we consider different haplotypes for simulations and inference but haplotypes are all trio-phased and admixture time is assumed to be known. The set D3 is the same as D2 except that admixture time is unknown and assume to be equal to 6 generations. In set D4, haplotypes used for inference are not true haplotypes but have been reconstructed with Beagle.

**Figure SI5:**
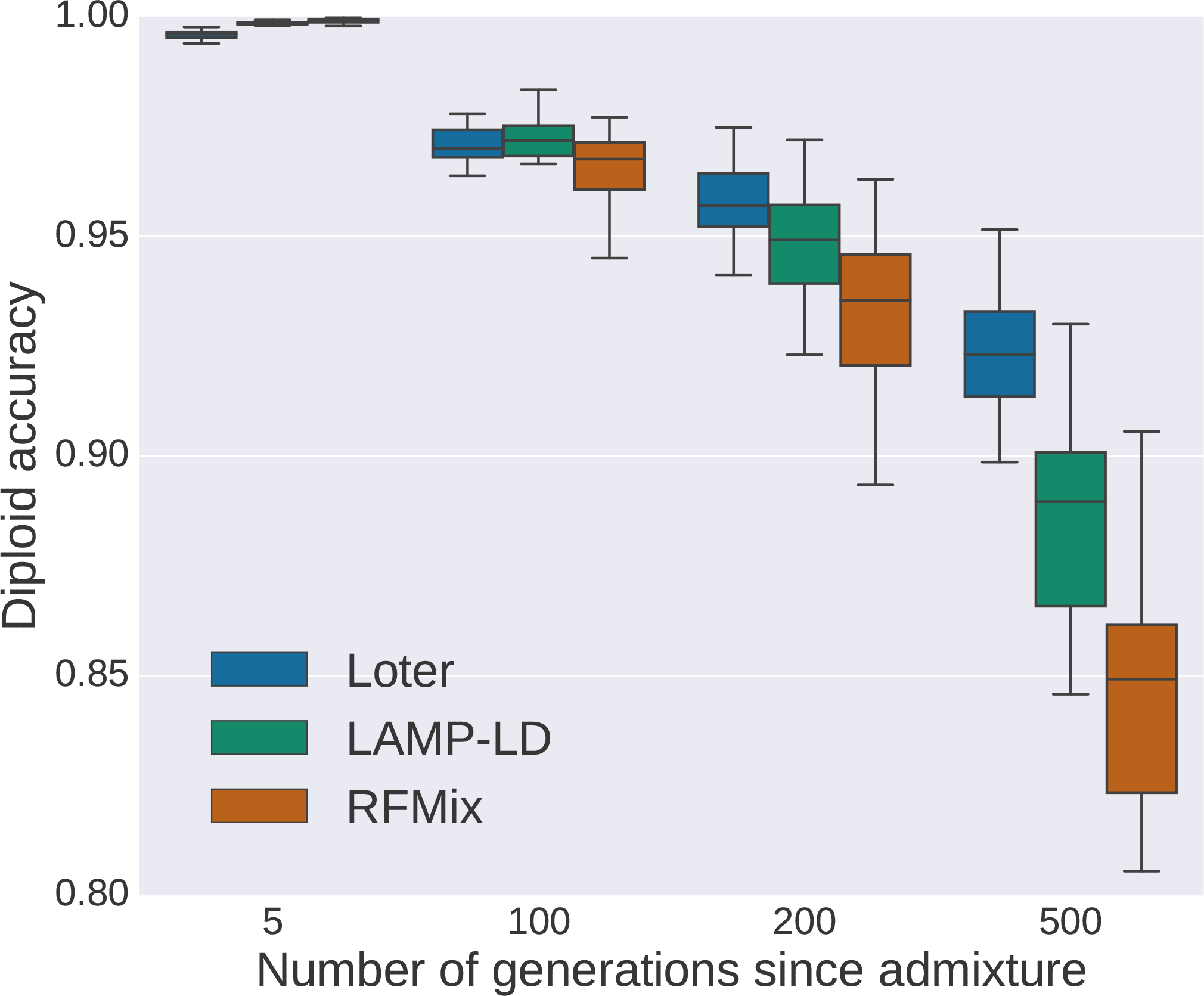
Diploid accuracy obtained under a 3-way admixture model with LAMP-LD, Loter, and RFMix for simulated admixed human individuals as a function of the time since admixture occurred. Admixed individuals are simulated by constructing their genomes from a mosaic of true African (YRI), European (CEU), and Chines haplotypes (International HapMap 3 Consortium et al. 2010). For performing simulations, true haplotypes are obtained using trio information. For local ancestry inference, haplotypes are also reconstructed using trio-based inference and are different from haplotypes used for simulations. For each value of the number of generations since admixture, 20 sets of 20 admixed individuals are generated. Boxplots show the distribution of the 20 values for the mean diploid accuracy.

**Figure SI6:**
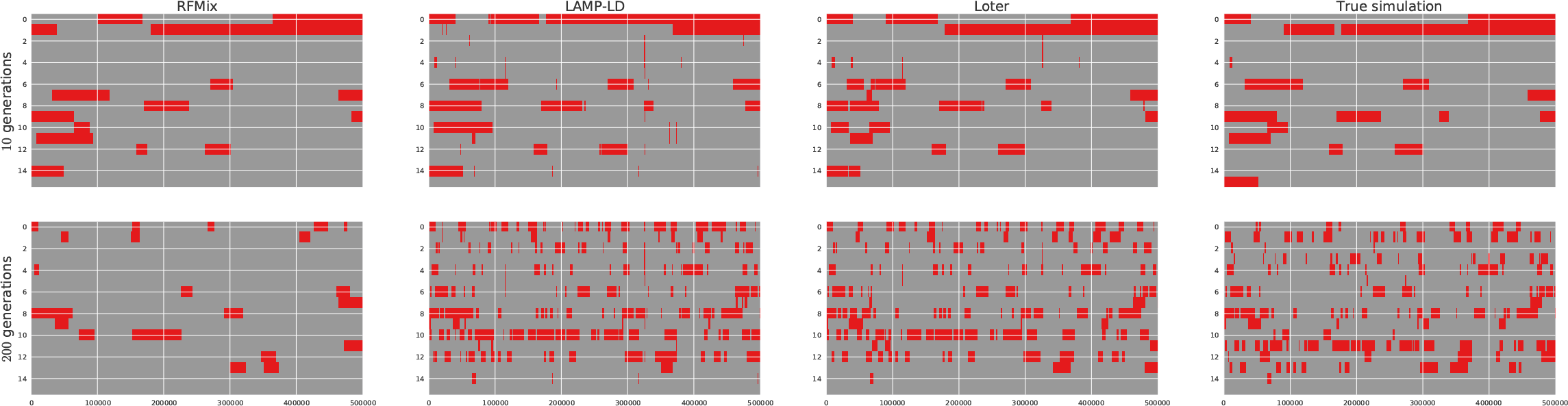
Ancestry tracts for 20 simulated admixed Populus individuals. Grey chunks correspond to *P. trichocarpa* chunks and red chunks correspond to *P. balsamifera* chunks. Two rows correspond to the two haplotypes of a single individual. Ancestry switches between haplotypes are caused by haplotype phasing using Beagle. The presence of spurious and small ancestry chunks contribute to excessively decrease the median length of ancestry chunks in LAMP-LD and Loter.

**Figure SI7:**
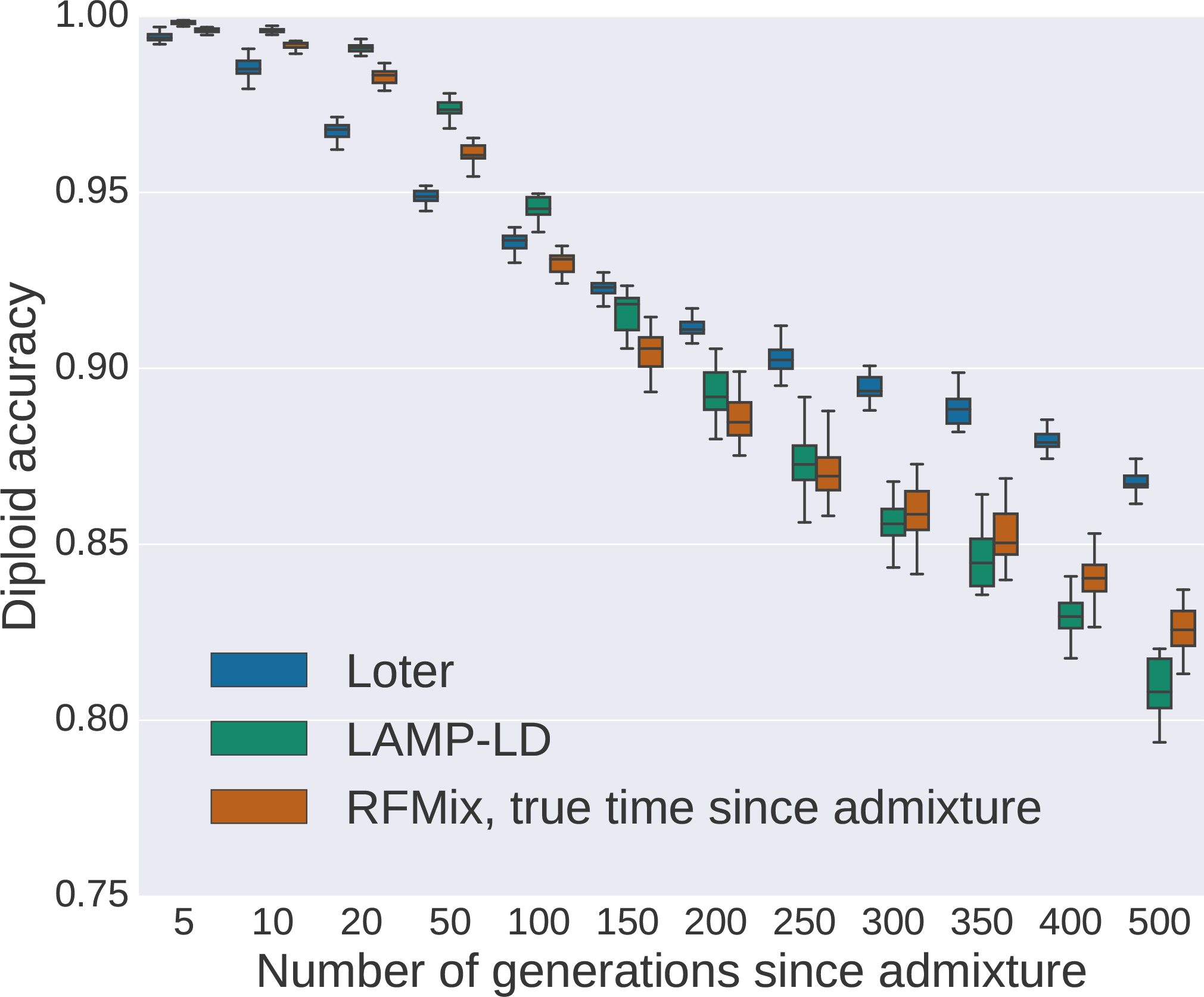
Diploid accuracy obtained with LAMP-LD, Loter, and RFMix for simulated admixed Populus individuals as a function of the time since admixture occurred when true values of the time since admixture are provided to RFMIX. Admixed individuals are simulated by constructing their genomes from a mosaic of *Populus trichocarpa* and *Populus balsamifera* individuals. Individuals are phased using Beagle and two different sets of individuals are used for performing simulations and inference. For each value of the number of generations since admixture, 20 sets of 20 admixed individuals are generated. Boxplots show the distribution of the 20 values for the mean diploid accuracy.

## References

Baran, Y., Pasaniuc, B., Sankararaman, S., Torgerson, D. G., Gignoux, C., Eng, C., Rodriguez-Cintron, W., Chapela, R., Ford, J. G., Avila, P. C. et al. 2012. Fast and accurate inference of local ancestry in Latino populations. Bioinformatics 28:1359–1367.

Bhatia, G., Patterson, N., Sankararaman, S. and Price, A. L. 2013. Estimating and interpreting F_ST_ : the impact of rare variants. Genome research 23:1514–1521.

Brandvain, Y., Kenney, A. M., Flagel, L., Coop, G. and Sweigart, A. L. 2014. Speciation and Introgression between *mimulus nasutus* and *mimulus guttatus*. PLoS Genet 10:e1004410.

Breiman, L. 1996. Bagging predictors. Machine learning 24:123–140.

Browning, S. R. and Browning, B. L. 2007. Rapid and accurate haplotype phasing and missing-data inference for whole-genome association studies by use of localized haplotype clustering. The American Journal of Human Genetics 81:1084–1097.

Browning, S. R. and Browning, B. L. 2011. Haplotype phasing: existing methods and new developments. Nature Reviews Genetics 12:703–714.

Bryc, K., Durand, E., Macpherson, J. M., Reich, D. and Mountain, J. 2015. The genetic ancestry of African, Latino, and European Americans across the United States. American Journal of Human Genetics 96:37–53.

Buerkle, C. A. and Lexer, C. 2008. Admixture as the basis for genetic mapping. Trends in ecology & evolution 23:686–694.

Corbett-Detig, R. and Nielsen, R. 2017. A hidden Markov model approach for simultaneously estimating local ancestry and admixture time using next generation sequence data in samples of arbitrary ploidy. PLoS genetics 13:e1006529.

Gravel, S. 2012. Population genetics models of local ancestry. Genetics 191:607–619.

Hufford, M. B., Lubinksy, P., Pyhäjärvi, T., Devengenzo, M. T., Ellstrand, N. C. and Ross-Ibarra, J. 2013. The genomic signature of crop-wild introgression in maize. PLoS Genet 9:e1003477.

International HapMap 3 Consortium et al. 2010. Integrating common and rare genetic variation in diverse human populations. Nature 467:52–58.

Kimmel, G., Sharan, R. and Shamir, R. 2003. Identifying blocks and sub-populations in noisy snp data. In International Workshop on Algorithms in Bioinformatics, 303–319. Springer.

Li, N. and Stephens, M. 2003. Modeling linkage disequilibrium and identifying recombination hotspots using single-nucleotide polymorphism data. Genetics 165:2213–2233.

Lindtke, D., Gonzalez-Martinez, S., Macaya-Sanz, D. and Lexer, C. 2013. Admixture mapping of quantitative traits in Populus hybrid zones: power and limitations. Heredity 111:474–485.

Liu, S., Lorenzen, E. D., Fumagalli, M., Li, B., Harris, K., Xiong, Z., Zhou, L., Korneliussen, T. S., Somel, M., Babbitt, C. et al. 2014. Population genomics reveal recent speciation and rapid evolutionary adaptation in polar bears. Cell 157:785–794.

Malinsky, M., Svardal, H., Tyers, A. M., Miska, E. A., Genner, M. J., Turner, G. F. and Durbin, R. 2017. Whole genome sequences of malawi cichlids reveal multiple radiations interconnected by gene flow. BioRxiv 143859.

Maples, B. K., Gravel, S., Kenny, E. E. and Bustamante, C. D. 2013. RFMix: a discriminative modeling approach for rapid and robust local-ancestry inference. The American Journal of Human Genetics 93:278–288.

Medugorac, I., Graf, A., Grohs, C., Rothammer, S., Zagdsuren, Y., Gladyr, E., Zinovieva, N., Barbieri, J., Seichter, D., Russ, I. et al. 2017. Whole-genome analysis of introgressive hybridization and characterization of the bovine legacy of Mongolian yaks. Nature Genetics 49:470–475.

Ni, X., Yang, X., Guo, W., Yuan, K., Zhou, Y., Ma, Z. and Xu, S. 2016. Length distribution of ancestral tracks under a general admixture model and its applications in population history inference. Scientific reports 6.

Paşaniuc, B., Sankararaman, S., Kimmel, G. and Halperin, E. 2009. Inference of locus-specific ancestry in closely related populations. Bioinformatics 25:i213–i221.

Patin, E., Siddle, K. J., Laval, G., Quach, H., Harmant, C., Becker, N., Froment, A., Régnault, B., Lemée, L., Gravel, S. et al. 2014. The impact of agricultural emergence on the genetic history of african rainforest hunter-gatherers and agriculturalists. Nature communications 5:3163.

Patterson, N., Hattangadi, N., Lane, B., Lohmueller, K. E., Hafler, D. A., Oksenberg, J. R., Hauser, S. L., Smith, M. W., O’Brien, S. J., Altshuler, D. et al. 2004. Methods for high-density admixture mapping of disease genes. The American Journal of Human Genetics 74:979–1000.

Payseur, B. A. and Rieseberg, L. H. 2016. A genomic perspective on hybridization and speciation. Molecular ecology 25:2337–2360.

Price, A. L., Tandon, A., Patterson, N., Barnes, K. C., Rafaels, N., Ruczinski, I., Beaty, T. H., Mathias, R., Reich, D. and Myers, S. 2009. Sensitive detection of chromosomal segments of distinct ancestry in admixed populations. PLoS Genet 5:e1000519.

Sankararaman, S., Kimmel, G., Halperin, E. and Jordan, M. I. 2008a. On the inference of ancestries in admixed populations. Genome research 18:668–675.

Sankararaman, S., Sridhar, S., Kimmel, G. and Halperin, E. 2008b. Estimating local ancestry in admixed populations. The American Journal of Human Genetics 82:290–303.

Scheet, P. and Stephens, M. 2006. A fast and flexible statistical model for large-scale population genotype data: applications to inferring missing genotypes and haplotypic phase. The American Journal of Human Genetics 78:629–644.

Seldin, M. F., Pasaniuc, B. and Price, A. L. 2011. New approaches to disease mapping in admixed populations. Nature Reviews Genetics 12:523–528.

Suarez-Gonzalez, A., Hefer, C. A., Christe, C., Corea, O., Lexer, C., Cronk, Q. C. and Douglas, C. J. 2016. Genomic and functional approaches reveal a case of adaptive introgression from *populus balsamifera* (balsam poplar) in *p. trichocarpa* (black cottonwood). Molecular ecology 25:2427–2442.

vonHoldt, B. M., Kays, R., Pollinger, J. P. and Wayne, R. K. 2016. Admixture mapping identifles introgressed genomic regions in North American canids. Molecular ecology 25:2443–2453.

Xue, J., Lencz, T., Darvasi, A., Pe’er, I. and Carmi, S. 2017. The time and place of european admixture in Ashkenazi Jewish history. PLoS genetics 13:e1006644.

Zhang, K., Deng, M., Chen, T., Waterman, M. S. and Sun, F. 2002. A dynamic programming algorithm for haplotype block partitioning. Proceedings of the National Academy of Sciences 99:7335–7339.

Zhou, Q., Zhao, L. and Guan, Y. 2016. Strong selection at MHC in Mexicans since admixture. PLoS genetics

